# Changes in Evidence Used for FDA Novel Drug Approvals Following the Implementation of the 21^st^ Century Cures Act

**DOI:** 10.64898/2026.01.31.702992

**Authors:** Aditya Narayan, Veronica L Irvin, Amanda J Koong, Sujin Song, Robert M Kaplan

## Abstract

**Background:** The 21st Century Cures Act (2017) expanded FDA flexibility in applying methodological standards for drug approval. To examine trends before and after implementation, we independently reviewed all novel drugs approved between 2016 and 2024.

**Methods:** We constructed a database of all novel FDA approvals from January 1, 2016, through December 31, 2024. Each study linked to an approved drug (N=6,763) was cataloged by study number, sponsorship, and timing of results reporting relative to completion.

**Results:** Since 2016, the number of studies supporting approval has steadily declined. Beginning in 2017, the modal number of studies per approval fell to one. Industry sponsorship increased while NIH-supported studies decreased. Average time to public posting of results exceeded the one-year statutory limit.

**Conclusions:** After implementation of the Cures Act, FDA approvals have relied on fewer, increasingly industry-sponsored studies. Although this may accelerate access to new therapies, it raises concerns about the strength of evidence for safety and effectiveness.

## Introduction

The U.S. Food and Drug Administration (FDA) serves a critical role in the regulation of medical technologies and pharmaceuticals, ensuring that new drugs and devices meet established standards for safety and efficacy before entering the market. The quality of evidence supporting drug approvals is of paramount importance. It underpins the FDA’s ability to make informed decisions that protect public health, ensuring that new treatments provide therapeutic value without introducing undue risks^1^. Maintaining high standards in the drug approval process is essential to prevent the introduction of therapies that may have adverse effects or insufficient efficacy^2^.

The 21st Century Cures Act, enacted in 2016, introduced changes to the FDA’s regulatory framework. The Act was intended to expedite the development and approval of medical products by promoting innovation and streamlining regulatory processes. Although the FDA previously permitted approval on the basis of a single pivotal trial when that trial is deemed adequate and well-controlled, the 21st Century Cures Act reinforced and broadened the agency’s flexibility to accept such evidence. In practice, the Act heightened concerns that approvals might increasingly rely on fewer pivotal studies, particularly in contexts where accelerated approval and surrogate endpoints are applied. This shift raised concerns within the medical and scientific communities regarding the potential implications for the robustness of the evidence supporting drug approvals.^3-8^ There is ongoing debate about whether these changes might compromise the thoroughness of the evaluation process, particularly in contexts where accelerated approval pathways and surrogate endpoints are utilized.^9-12^

The implications of sponsorship and funding in drug approval processes are critical to understanding the potential biases and influences that may affect the integrity of clinical trials. Industry funding, which has become increasingly dominant, raises concerns about the objectivity of research outcomes, as sponsors may have vested interests in favorable results that expedite the approval of drugs. This potential conflict of interest can lead to the prioritization of speed over thoroughness in the evaluation process, particularly under frameworks like those established under the 21st Century Cures Act, which allows for accelerated approvals. Additionally, the diversification of funding sources, including collaborations between industry, government agencies, and academic institutions, introduces complexities in ensuring the transparency and accountability of trial data.

Further, evaluating the reporting of trial results is crucial for ensuring transparency and accountability in the drug approval process^13-15^. The 2017 Food and Drug Administration Amendments Act (FDAAA) introduced penalties for failing to report trial results within one year of study completion. Despite the threat of fines up to $10,000 per day, with a potential ceiling of $1 million, the policy has had limited impact since its implementation in 2017. A key reason for this is the lack of enforcement by the FDA. By 2021, four years after the policy’s introduction, the FDA had issued only four notices of noncompliance and had not collected any fines. Uncollected potential fines were estimated to total $19 billion^13^—a sum greater than five times the FY 2025 congressional appropriation for the agency. This lack of enforcement has resulted in a continued pattern of underreporting, potentially compromising the integrity of the drug approval process^14,15^.

Now, eight years following the implementation of the 21^st^ Century Cures Act, Congress has begun to consider the next generation of the initiative (Cures 2.0)^16^. The proposed Cures 2.0 Act is designed to build on the successes of the original by accelerating medical innovation, expanding access to telehealth, and modernizing the U.S. healthcare system while minimizing its unintended consequences. Key provisions of the bill include enhancing diversity in clinical trials, expanding the use of real-world evidence (RWE) to fulfill post-marketing commitments, and improving the inclusion of patient experience data. Additionally, the legislation seeks to increase Medicare coverage for new health technologies and expand access to telehealth services by removing geographic restrictions.

In light of these proposed changes, understanding the trends in FDA trial approvals and funding sources over recent years becomes crucial. This study examines trends in the number of clinical trials utilized in FDA approvals from 2016 to 2024 with the objective of determining whether there has been a change in evidentiary standards over this period^17^. Such context will be necessary for evaluating whether policies under Cures 2.0 might further shift standards – particularly as real-world evidence and accelerated approvals become more prominent – and, in turn, shape informed policy decisions aimed at safeguarding both innovation and patient safety^18^.

## Methods

We conducted a retrospective review of all Novel Drug Approvals (NDAs) by the U.S. Food and Drug Administration (FDA) from 2016 to 2024. Data on each approved drug were obtained from the FDA’s publicly accessible records, including the number of trials in the FDA report. To gather detailed information on the studies associated with these approvals, we used the National Library of Medicine’s ClinicalTrials.gov database. For each approved medication, we extracted data on the timing and characteristics of each associated study, including the date when each study was first posted, the start date, the primary completion date, and the date when results were reported, if applicable. Additional variables included study design and sponsorship. Sponsorship categories were coded hierarchically: trials were classified as “industry” if industry support was present, even when combined with other sources. Joint industry-government or NIH-other sponsorships were recorded separately to avoid misclassification. All coders collected variables for a subset of approved drugs each year to ensure standardization. We systematically downloaded all standardized fields from ClinicalTrials.gov into a comprehensive dataset for further analysis.

### Completed Trials

For each drug approval, we identified and coded the total number of studies registered, the number of trials used directly in the FDA approval process —as identified in FDA approval packages—or closely related studies cited in the approval rationale, and the number of completed studies. A study was classified as completed if the primary completion date occurred before the drug’s approval date. We repeated the above calculations separately for studies with randomized controlled designs.

### Analysis

Descriptive statistics, including mean values and standard deviations, were calculated for variables such as the number of trials per approval. All analyses were performed using R version 4.2.0. A Pearson chi-square test was employed to evaluate changes in the distribution of the number of trials used for approval (categorized as one, two, or three or more trials) across the years, with statistical significance set at p < 0.05. Additionally, we used Analysis of Variance (ANOVA) to assess linear trends in the mean number of trials over time. Crosstabulations and bar plots were utilized to examine shifts in funding sources for studies throughout the observed period. The difference between the date of primary completion of each study (the date that the last data point for the primary outcome measure was collected from the last enrolled participant) and the date of results posting was used to identify delays in reporting. Studies with missing data for result posting dates or primary completion dates were excluded from the analysis.

## RESULTS

### Number of Trials in Approvals

Between 2016 and 2024, there was a linear decline in the mean number of studies used per approval (Figure 1). In 2016, the mean number of studies was 3.41, which decreased to 3.11 in 2017. This figure continued to decline to 1.50 by 2022. However, in 2023, there was an increase to a mean of 2.49 trials per approval. This was attributable to a single outlier (Brenzavvy; an SGLT2i), which was approved based on nine trials. Without this outlier, the mean number of trials used for approval in 2023 was 1.85. The linear trend toward fewer trials in the approval by year continued in 2024, when the mean number of trials in the approval was 1.39. An Analysis of Variance planned contrast for linear trend was highly significant (F _1/6753_=417.98,p<.0001).

**Figure 1.**
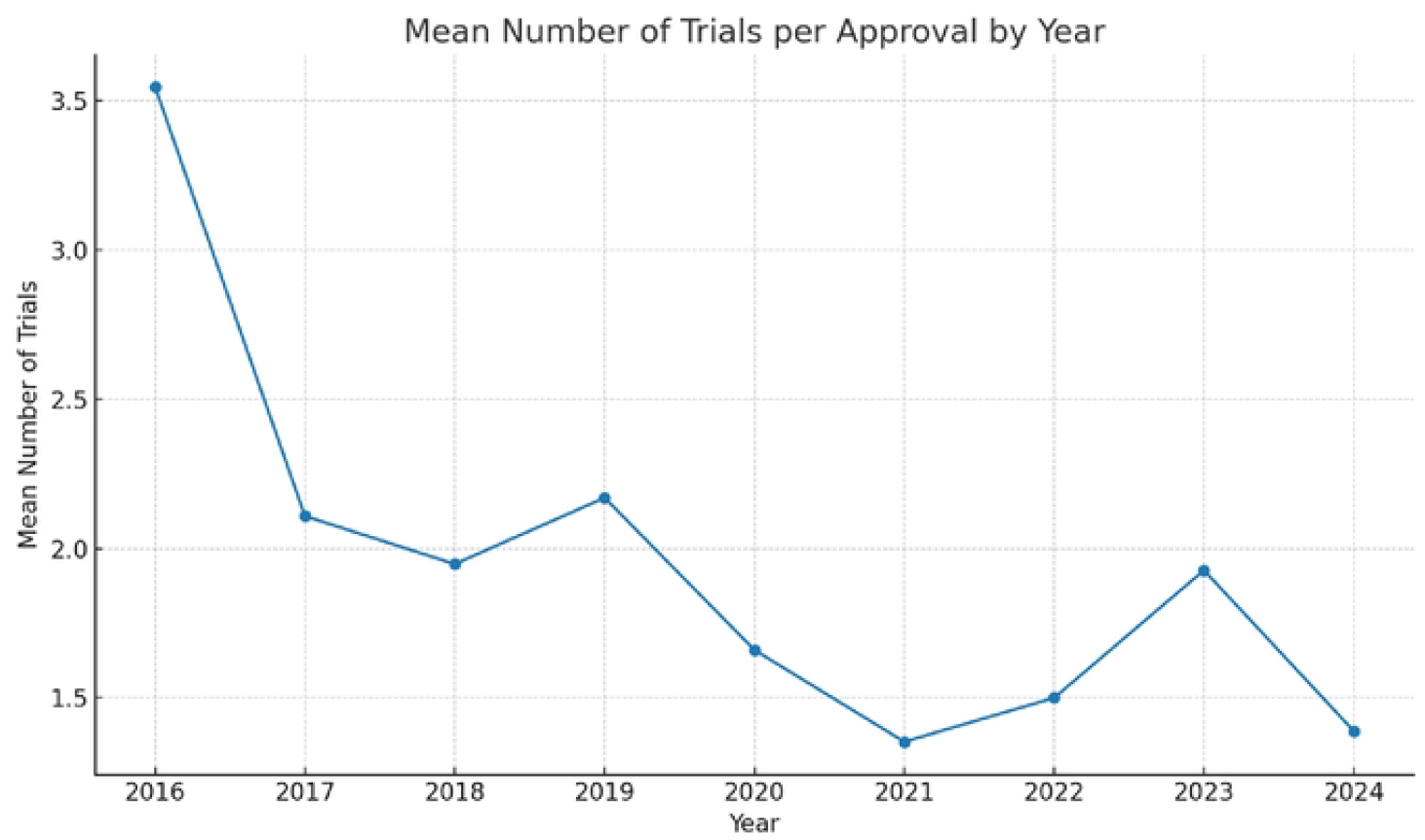
Mean number of trials in FDA approval by year

In addition to the mean number of trials in approvals, we examined the trend in the number of approvals supported by three or more trials. Notably, in 2016, the majority of drugs (59.1%) were approved based on three or more trials. However, by 2021, this proportion had significantly decreased, with only 11.8% of drugs being approved based on three or more trials. Instead, the majority of approvals in 2021 (76.5%) were based on a single trial. By 2024, 69.4% of products were approved on the basis of a single trial, and only 5.6% were approved on the basis of 3 or more trials. The Pearson chi-square test of these findings indicates that the distribution of the number of trials used in drug approvals has changed significantly over the years studied (X^2^ = 48.19; p < 0.001)(Figure 2), with a significant linear decline in the number approved on the basis of 3 or more studies X^2^ = 26.20, df =8, p<.0001) The Analysis of Variance contrast test for linear trend was highly significant (F _1/6006_ = 129.96, p<.001) (Figure 2).

**Figure 2.**
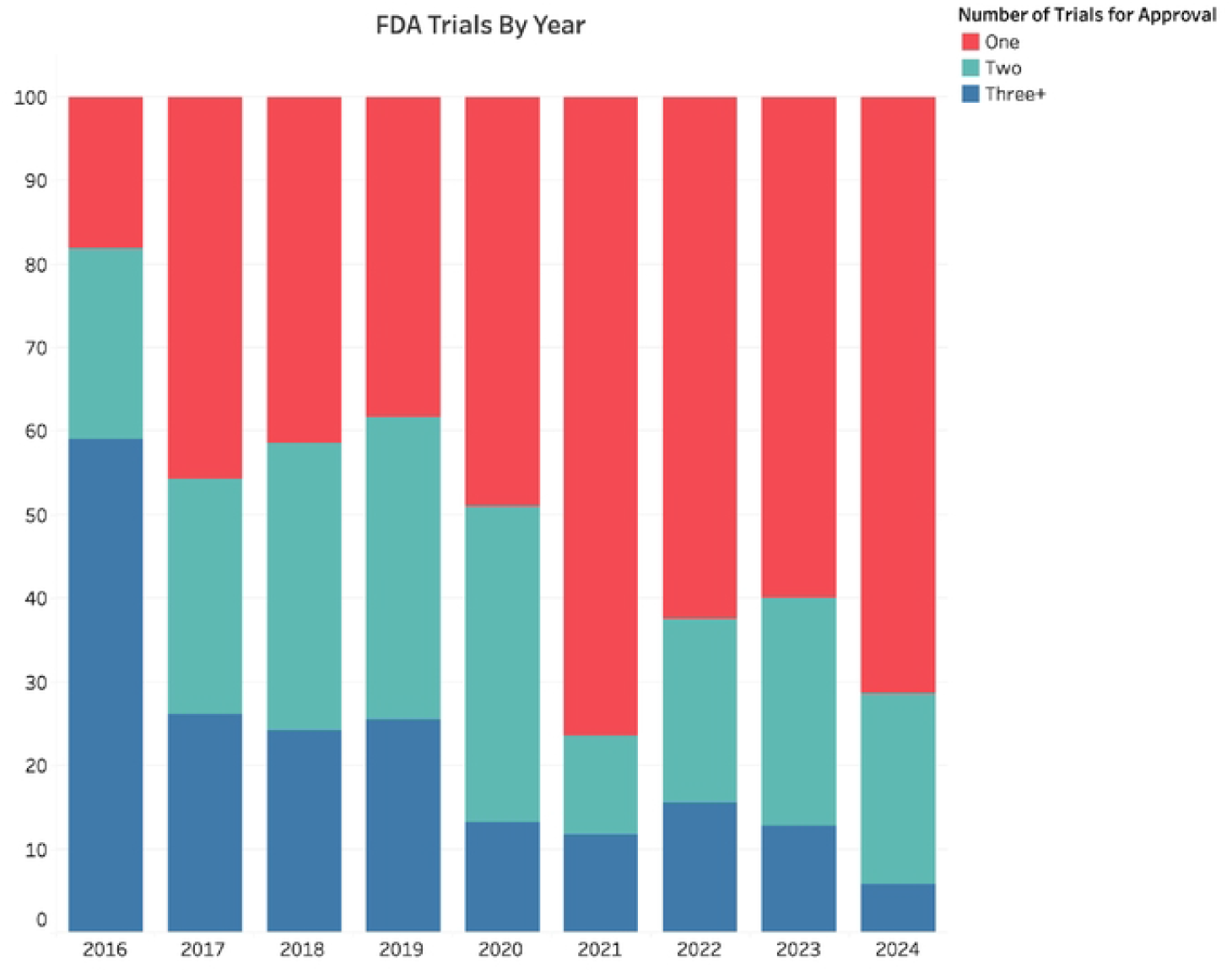
Approvals with one, two, or three or more trials used in FDA approval: 2016-2023.

### Funding Sources for Studies

Across the observed years, industry funding consistently dominated study sponsorship, with a general upward trend in its share of total funding (Table 1). In 2016, industry-funded trials accounted for 44.6%, while non-industry sources accounted for 55.4%. Over time, the industry’s share steadily increased, reaching 55.2% in 2018 and climbing to 66.4% by 2019 and 80.4% by 2024. During this period, NIH and other government sources played a relatively minor role, contributing less than 3% of total study funding in most years. Regression analysis showed a significant linear trend over years (percent industry funded =0.037 X year – 73.74; R^2^=.45, p<.01).

**Table 1.** Proportion of sponsor types for studies registered in ClinicalTrials.gov by the year of FDA approval for that drug.

|  | 2016 | 2017 | 2018 | 2019 | 2020 | 2021 | 2022 | 2023 | 2024 |
| --- | --- | --- | --- | --- | --- | --- | --- | --- | --- |
| Industry | 44.6% | 43.5% | 55.2% | 66.4% | 42.2% | 68.5% | 66.1% | 69.9% | 80.4% |
| Industry & Other | 0% | 12.1% | 0% | 0% | 0% | 10.5% | 8.4% | 0% | 0% |
| NIH | 3% | 0.7% | 2.3% | 1.3% | 3.6% | 2.3% | 0.5% | 0.4% | 0.3% |
| NIH & Other | 0% | 2.4% | 0% | 0% | 0% | 2.9% | 3.1% | 0% | 0% |
| Other | 52.4% | 41.3% | 42.5% | 32.3% | 54.2% | 15.7% | 22% | 29.7% | 19.3% |

### Timing of Study Results Reporting

In addition to examining evidentiary standards, we also assessed the timeliness of results reporting as a secondary outcome, given its importance for transparency. The duration between a study’s primary completion date and the public posting of its results exhibited considerable variability from 2016 to 2023 (Figure 3). Clinical trials demonstrated a similar trend, with the average days to result posting starting at 840.7 days in 2016 and increasing to 1,226 days in 2020. This period was followed by a reduction and subsequent increase. Accordingly, while there were periods of both extension and reduction in the time taken to post results, the average time remained consistently above one year for studies with available results throughout the observed years. The average time to result posting was consistently longer for clinical trials compared to all trials, highlighting differences in reporting timelines between these study types.

**Figure 3.**
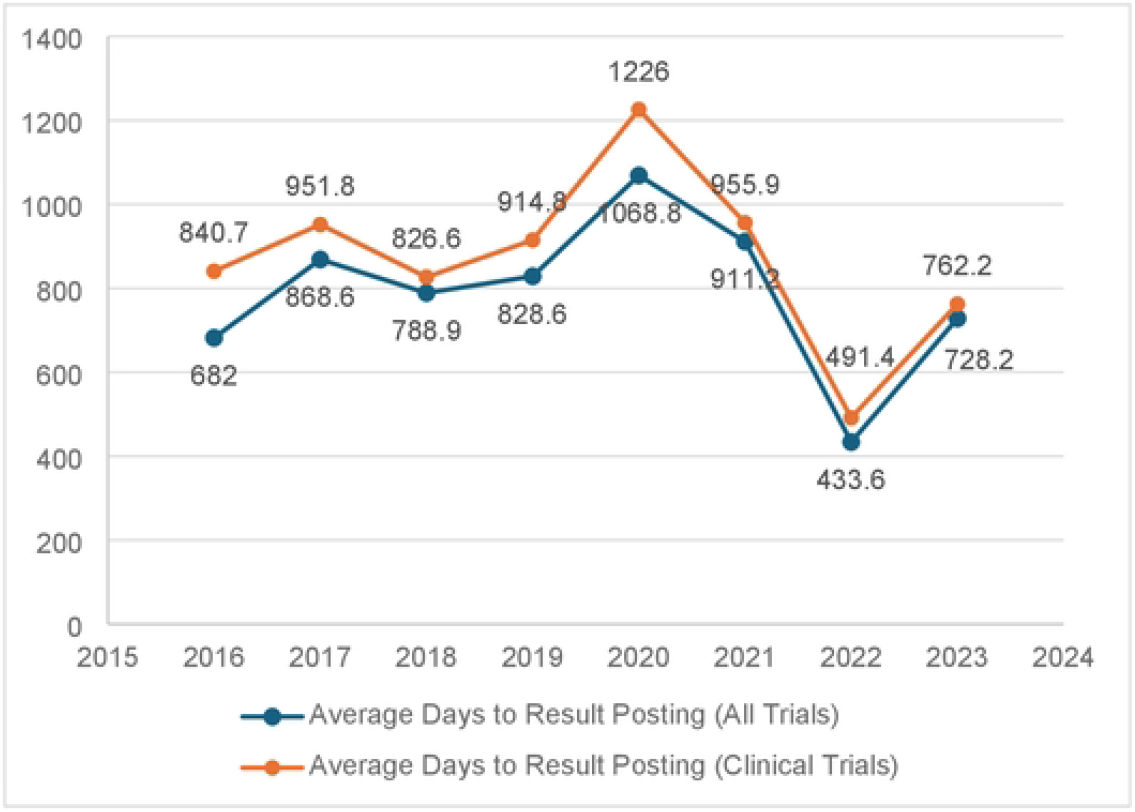
Average Days from Primary Completion Date to Result Reporting. Clinical trials were defined as randomized interventional studies as indicated on ClinicalTrials.Gov.

## Discussion

Since the 2017 implementation of the 21^st^ Century Cures Act, the number of FDA novel drug approvals has increased^19^ while the number of trials used to justify each new approval has systematically declined^11^. Our observation of a decrease in the number of trials used for FDA drug approvals from 2017 to 2024 raises important considerations for the regulatory process and drug development strategies. The FDA now appears to focus on fewer clinical trials to assess each drug’s efficacy and safety^20^. Additionally, our data reveal consistent delays in the reporting of clinical trial results, with average times exceeding one year. These prolonged reporting delays may hinder the timely evaluation of drug safety and efficacy, potentially impacting the regulatory decision-making process and reducing the transparency of the approval pathway. The combination of reduced trial numbers and extended result reporting timelines underscores the need for enhanced regulatory oversight to ensure that the evidence base supporting drug approvals remains robust and reliable.

Although we cannot assure the relationship is causal, the trend in approvals coincided with federal policy changes. The implementation of the 21st Century Cures Act, which allows for greater flexibility in drug approvals, facilitated approvals based on data from a single pivotal trial instead of multiple well-controlled studies. While this administrative change allowed earlier access to new medications, it may have led to approvals based on less comprehensive evidence. More rapid dissemination also raised concerns about the long-term safety and effectiveness of drugs that become available to consumers^21^.

Of note, the majority of pivotal evidence continued to come from phase 3 trials across all years studied. While the absolute number of phase 3 trials per approval declined in parallel with the overall decrease in trials, we did not observe a compensatory increase in reliance on earlier phase (phase 1 or 2) studies as the primary basis for approval. This suggests that the trend reflects fewer late-stage trials being conducted or submitted per approval, rather than a systematic substitution of early-stage evidence for confirmatory studies

Building on concerns related to the reduced number of trials used in FDA approvals, we examined the funding landscape of clinical trials over the same period. A review of FDA approvals in 2022 showed that 65% of drugs were approved based on a single study, a substantial increase from previous years^22^. Our analysis of funder types for FDA drug approvals from 2016 to 2023 reveals a clear and consistent trend: the increasing dominance of industry funding in studies associated with drug approvals. Industry sponsors accounted for the majority of funding across all years while funding from other sources, including the National Institutes of Health (NIH) and government networks, remained minimal and even declined over time. For instance, the proportion of trials with NIH funding decreased from 2.97% in 2016 to just 0.41% in 2023 (while the NIH’s overall budget has remained level or increased over this time frame). Our observations confirm analogous trends reported by Buchkowsky^23^ and others^23,24^. This shift in funding sources has significant implications for the types of evidence presented in support of new drug approvals.

The increasing reliance on industry funding raises concerns about potential conflicts of interest and the influence of sponsors on the design, conduct, and reporting of clinical trials^25^. Research has shown that industry-sponsored trials are more likely to yield favorable outcomes for sponsors’ products, which can influence the approval process and potentially compromise the safety and efficacy of approved drugs^26,27^. The potential for selective reporting and publication bias is heightened when industry sponsors control the narrative around a drug’s efficacy and safety, as they may prioritize the dissemination of favorable results while minimizing or withholding less favorable data.

The implications of this trend for public policy, particularly in light of the proposed Cures 2.0 initiative, are significant. As industry funding becomes more dominant, while journals and ClinicalTrials.gov do mandate sponsor disclosure and author conflicts of interest, there may be a need for greater transparency and accountability in the drug approval process. The FDA may need to revisit its approval criteria to ensure that the increasing reliance on industry-funded trials does not compromise the safety and efficacy of approved drugs^28^. This could involve strengthening requirements for independent replication of industry-sponsored studies or mandating the inclusion of data from trials funded by non-industry sources, such as the NIH, in the approval process. Maintaining public trust in the FDA’s regulatory process is essential, and this can only be achieved by ensuring that drug approvals are based on a comprehensive and unbiased evaluation of all available evidence.

Our analysis also raises concerns about the completion and timing of result reporting in clinical trials, with an evaluation of the timing of result reporting revealing that the average duration between a study’s primary completion date and the public posting of its results consistently exceeded one year throughout the study period (Figure 3). Notably, there were significant fluctuations during the COVID-19 pandemic years of 2020 and 2021. In 2020, the average days to result posting peaked across all study years at 1,068.8 days for all trials and 1,226 days for clinical trials, reflecting substantial delays. These delays may be attributable to the unprecedented pressures and resource reallocations faced by the FDA and research institutions during the pandemic, as well as the expedited approval processes for critical therapies under Emergency Use Authorizations (EUAs). As a result, the FDA’s capacity to monitor and enforce standard result reporting deadlines for non-COVID-related trials may have been strained. The prioritization of pandemic-related approvals could inadvertently lead to delays in the oversight and timely reporting of results for other ongoing studies. Additionally, while this flexibility is essential for addressing urgent public health needs, it can also mean that certain administrative processes, including result reporting, are deprioritized.

Although there was a noticeable reduction in average days to result posting in 2022 – potentially reflecting a return to more typical approval processes – the trend underscores persistent challenges in achieving timely transparency. To this end, more recently, the Food and Drug Omnibus Reform Act (FDORA) of 2022 has further emphasized reforms to accelerated approval, including requirements for confirmatory studies to be underway at the time of approval. These changes may help address some of the concerns identified in our analysis.^29^

Our finding of long delays between trial completion and the public posting of results on ClinicalTrials.gov is consistent with other reports in the published literature ^30^. Furthermore, the completeness of reporting, particularly regarding adverse events, is often more comprehensive in ClinicalTrials.gov than in corresponding journal articles, underscoring the importance of timely and thorough data dissemination ^30^. The variability in trial completion and reporting times also raises concerns about the robustness of the evidence supporting FDA approvals, particularly for drugs approved under accelerated pathways^31^. Delayed or incomplete reporting can hinder independent verification of trial outcomes, which is critical for ensuring that approved drugs meet the necessary safety and efficacy standards. Accordingly, policies could be strengthened by increasing penalties for non-compliance, mandating real-world data collection post-approval, and improving transparency in trial result disclosures. Given the evolving FDA and HHS leadership, federal priorities have the potential to shape how flexibly existing pathways are interpreted, and how strongly post-approval commitments are enforced.

## Limitations

The results from our study should be interpreted in light of several limitations. First, the FDA only reports information on approved drugs, providing limited public information about products that were evaluated but not approved. Similarly, a substantial number of new molecular entities are abandoned based on null or negative trial findings. In these cases, the regulatory system may be functioning as intended by discouraging the continued evaluation of drugs with low efficacy or high safety concerns. Second, our analysis is dependent on the quality of information in ClinicalTrials.gov. Although the database is overseen by experts at the National Library of Medicine, entries are not peer-reviewed. It is likely that the information database has not undergone the same level of scrutiny that characterizes evaluations of NIH grants and journal publications. Third, we collected all of the data within a relatively short time period, resulting in differing lengths of follow-up for studies associated with drug approvals from 2016 to 2024. Specifically, drugs approved in 2016 had up to seven years for associated studies to be registered and reported, whereas data for drugs approved in 2024 were collected at the end of 2024 or the beginning of 2025. This discrepancy may lead to incomplete registration and reporting of studies for more recent drug approvals, potentially underestimating the number of trials and affecting the analysis of trial utilization over time. Fourth, the presence of outliers, such as the Brenzavvy (an SGLT2i) approved based on nine trials in 2023, may skew the mean number of trials used per approval, although sensitivity analyses were conducted to assess their impact. Fifth, the concept of ‘completion’ differs substantially across trials, with laboratory-based surrogate endpoints often requiring shorter follow-up than trials measuring long-term clinical outcomes. This heterogeneity likely contributes to observed variation in trial duration and reporting. These limitations highlight the need for cautious interpretation of our findings and suggest areas for future research to enhance the robustness and comprehensiveness of data on FDA drug approvals.

## Conclusions

An independent review of all FDA novel drug approvals between 2016 and 2024 identified significant shifts in the FDA drug approval process, particularly in the decreasing number of trials used in approval and the increasing reliance on industry-funded studies. Additionally, the consistent delays in trial result reporting highlight the need for improved mechanisms to ensure timely dissemination of clinical data. These trends point to the need for policy adjustments, including stronger enforcement of trial reporting requirements, greater transparency, and increased involvement of non-industry funding sources to mitigate potential biases. Moving forward, policies that bolster post-market surveillance, enforce timely and comprehensive trial result reporting, and support independent clinical research are crucial for maintaining the integrity of the drug approval process^32^.

## References

1. Kowitt SD, Schmidt AM, Hannan A, Goldstein AO. Awareness and trust of the FDA and CDC: Results from a national sample of US adults and adolescents. PloS one. 2017;12(5):e0177546.

2. Brown BL, Mitra-Majumdar M, Joyce K, et al. Trends in the quality of evidence supporting FDA drug approvals: results from a literature review. Journal of Health Politics, Policy and Law. 2022;47(6):649–672.

3. Mendoza RL. The 21st Century Cures Act: pharmacoeconomic boon or bane? : Taylor & Francis; 2017. p. 315–317.

4. Sarpatwari A, Kesselheim A. The 21st century cures act: opportunities and challenges. Clinical Pharmacology & Therapeutics. 2015;98(6):575–577.

5. Goble JA. The potential effect of the 21st century cures act on drug development. Journal of Managed Care & Specialty Pharmacy. 2018;24(7):677–681.

6. Kesselheim AS, Avorn J. New “21st Century Cures” legislation: speed and ease vs science. Jama. 2017;317(6):581–582.

7. Hudson KL, Collins FS. The 21st Century Cures Act—a view from the NIH. New England Journal of Medicine. 2017;376(2):111–113.

8. GOV O. How FDA Used Its Accelerated Approval Pathway Raised Concerns in 3 of 24 Drugs Reviewed.

9. Darrow JJ, Avorn J, Kesselheim AS. The FDA breakthrough-drug designation— four years of experience. N Engl J Med. 2018;378(15):1444–1453.

10. Darrow JJ, Avorn J, Kesselheim AS. FDA Approval and Regulation of Pharmaceuticals, 1983-2018. JAMA. 2020;323(2):164–176.

11. Kaplan RM, Koong AJ, Irvin V. Review of Evidence Supporting 2022 US Food and Drug Administration Drug Approvals. JAMA Network Open. 2023;6(8):e2327650–e2327650.

12. Kaplan RM, Koong AJ, Irvin V. Food and Drug Administration novel drug decisions in 2017: transparency and disclosure prior to and 5 years following approval. Health Affairs Scholar. 2023;1(2):qxad028.

13. Stephenson J. In a first, FDA warns company to remedy failure to post clinical trial results. American Medical Association; 2021:e211306–e211306.

14. Piller C. FDA and NIH let clinical trial sponsors keep results secret and break the law. Science. 2020;10

15. DeVito NJ, Bacon S, Goldacre B. Compliance with legal requirement to report clinical trial results on ClinicalTrials. gov: a cohort study. The Lancet. 2020;395(10221):361–369.

16. Reps. DeGette, Bucshon Seek Stakeholder Input on Next-Generation Cures Bil. 2024. https://degette.house.gov/media-center/press-releases/reps-degette-bucshon-seek-stakeholder-input-next-generation-cures-bill

17. Nelson JT, Tse T, Puplampu-Dove Y, Golfinopoulos E, Zarin DA. Comparison of availability of trial results in ClinicalTrials. gov and PubMed by data source and funder type. JAMA. 2023;329(16):1404–1406.

18. Kadakia KT, Krumholz HM. Designing cures 2.0—from corridors to cornerstones. New England Journal of Medicine. 2022;386(18):1677–1679.

19. Brown DG, Wobst HJ. A decade of FDA-approved drugs (2010–2019): trends and future directions. Journal of medicinal chemistry. 2021;64(5):2312–2338.

20. Reardon S. FDA approves Alzheimer’s drug amid safety concerns. Nature. 2023;613(7943):277–278.

21. Zuckerman D, Jury N, Silcox C. 21st Century Cures Act: fast-track to cure or morbidity. Pharm Med. 2016;30:117–118.

22. de la Torre BG, Albericio F. The pharmaceutical industry in 2022: an analysis of FDA drug approvals from the perspective of molecules. Molecules. 2023;28(3):1038.

23. Buchkowsky SS, Jewesson PJ. Industry sponsorship and authorship of clinical trials over 20 years. Annals of Pharmacotherapy. 2004;38(4):579–585.

24. Flacco ME, Manzoli L, Boccia S, et al. Head-to-head randomized trials are mostly industry sponsored and almost always favor the industry sponsor. Journal of clinical epidemiology. 2015;68(7):811–820.

25. Lexchin J, Bero LA, Djulbegovic B, Clark O. Pharmaceutical industry sponsorship and research outcome and quality: systematic review. bmj. 2003;326(7400):1167–1170.

26. Lundh A, Lexchin J, Mintzes B, Schroll JB, Bero L. Industry sponsorship and research outcome. Cochrane database of systematic reviews. 2017;(2)

27. Mitra-Majumdar M, Gunter SJ, Kesselheim AS, et al. Analysis of supportive evidence for US Food and Drug Administration approvals of novel drugs in 2020. JAMA Network Open. 2022;5(5):e2212454–e2212454.

28. Karas L. FDA’s revolving door: reckoning and reform. Stan L & Pol’y Rev. 2023;34:1.

29. Narayan A, Cohen IG, Adashi EY. Food and Drug Omnibus Reform Act: a critical course correction. Elsevier; 2024:869–872.

30. Riveros C, Dechartres A, Perrodeau E, Haneef R, Boutron I, Ravaud P. Timing and completeness of trial results posted at ClinicalTrials. gov and published in journals. PLoS medicine. 2013;10(12):e1001566.

31. Diu S. Slowing Down Accelerated Approval: Examining the Role of Industry Influence, Patient Advocacy Organizations, and Political Pressure on FDA Drug Approval. Fordham L Rev. 2021;90:2303.

32. Zarin DA, Tse T, Williams RJ, Califf RM, Ide NC. The ClinicalTrials. gov results database—update and key issues. New England Journal of Medicine. 2011;364(9):852–860.

